# MHCSeqNet: A deep neural network model for universal MHC binding prediction

**DOI:** 10.1101/371591

**Authors:** Poomarin Phloyphisut, Natapol Pornputtanapong, Sira Sriswasdi, Ekapol Chuangsuwanich

**Affiliations:** Department of Computer Engineering, Faculty of Engineering, Chulalongkorn University, 254 Phayathai Road, Pathumwan, Bangkok 10330, Thailand; Department of Biochemistry and Microbiology, Faculty of Pharmaceutical Sciences, Chulalongkorn University, 254 Phayathai Road, Pathumwan, Bangkok 10330, Thailand; Vaccine and Therapeutic Protein, the Special Task Force for Activating Research, Faculty of Pharmaceutical Sciences, Chulalongkorn University, Bangkok 10330, Thailand; Computational Molecular Biology Group, Faculty of Medicine, Chulalongkorn University, 1873 Rama IV Road, Pathum Wan, Bangkok 10330, Thailand; Research Affairs, Faculty of Medicine, Chulalongkorn University. 1873 Rama IV Road, Pathum Wan, Bangkok 10330, Thailand

**Author notes:** To whom the correspondence should be addressed. Ekapol Chuangsuwanich. Natapol Pornputtanapong. Sira Sriswasdi.

## Abstract

Immunotherapy is an emerging approach in cancer treatment that activates the host immune system to destroy cancer cells that express unique peptide signatures (neoepitopes). Administrations of cancer-specific neoepitopes in the form of synthetic peptide vaccine have been proven effective in both mouse models and human patients. Because only a tiny fraction of cancer-specific neoepitopes actually elicit immune response, selection of potent, immunogenic neoepitopes remains a challenging step in cancer vaccine development. A basic approach for immunogenicity prediction is based on the premise that effective neoepitope should bind with the Major Histocompatibility Complex (MHC) with high affinity. In this study, we developed MHCSeqNet, an open-source deep learning model, which not only competes favorably with state-of-the-art predictors but also exhibits promising generalization to new MHC class I and class II alleles. MHCSeqNet employed neural network architectures developed for natural language processing to model amino acid sequence representations of MHC allele and epitope peptide as sentences with amino acids as individual words. This consideration allows MHCSeqNet to accept new MHC alleles as well as peptides of any length. The flexibility offered by MHCSeqNet should make it a valuable tool for screening effective neoepitopes in cancer vaccine development.

**Availability and implementation:** https://github.com/cmbcu/MHCSeqNet

## INTRODUCTION

Immunotherapy is a promising approach in cancer treatment that activates the host immune system to specifically destroy cancer cells, with far fewer adverse effects than chemotherapy or radiotherapy. This is possible because cancer cells produce unique peptide signatures (neoepitopes) derived from mutated proteins or normal proteins with tumor-specific expression, some of which are presented on the cancer cells’ outer surface and recognized by T cells (Castle *et al.*, 2012; Schumacher and Schreiber, 2015). Administrations of vaccines composed of synthetic peptides resembling cancer-specific neoepitopes have been proven to boost T cell activity to destroy cancer cells in both mouse models and human patients (Banchereau and Palucka, 2017; Sahin and Türeci, 2018; Schumacher and Schreiber, 2015). Nonetheless, because only a tiny fraction of hundreds of cancer-specific neoepitopes can elicit immune response, selection of effective immunogenic neoepitopes remains a challenging step in cancer vaccine development.

A basic approach for immunogenicity prediction is based on the fact that the Major Histocompatibility Complex (MHC), also called Human Leukocyte Antigen (HLA) complex, binds to peptide epitopes and presents them on the outer cell surface for recognition by T cells. In other words, a good neoepitope should be able to bind with MHC molecule with high affinity (Engels *et al.*, 2013). Current state-of-the-art software tools for peptide-MHC binding affinity prediction, such as NetMHCPan (Jurtz *et al.*, 2017), were trained using a combination of binding affinity data obtained from laboratory experiments and eluted ligand data from mass spectrometry analyses (Abelin *et al.*, 2017; Vita *et al.*, 2015) and exhibited high accuracy. Although recent approaches that employed deep learning models have demonstrated performance gains over NetMHCPan - the established tool-of-choice in many cancer vaccine development pipelines, they lacked NetMHCPan’s capability to make prediction for peptides of any lengths and for new MHC alleles that are not present in the training dataset.

In this study, we developed MHCSeqNet, an open-source deep learning model that can predict peptide-MHC binding affinity with high accuracy and with no restriction on the length of the peptide or the MHC allele, as long as its amino acid sequence is known. Our training and testing datasets were derived from the Immune Epitope Database (IEDB) (Vita *et al.*, 2015) and recent publications (Bassani-Sternberg *et al.*, 2016; O’Donnell *et al.*, 2018). The pooled dataset was cleaned by removing inconsistent and low-confidence entries.

Amongst the different fundamental architectures for deep learning, Recurrent Neural Networks (RNNs) have been used to encode time series information in many tasks such as automatic speech recognition (Graves *et al.*, 2013), natural language processing (Bahdanau *et al.*, 2014), and also bioinformatics (Quang and Xie, 2016). Unlike fully connected feed forward networks, RNNs can better capture temporal relationships by remembering previous inputs. A popular choice for RNNs is the Long Short-Term Memory (LSTM) which alleviates the vanishing and exploding gradient problems presented in normal RNNs. Gated Recurrent Unit (GRU) is a recently developed model which can be considered as a simplified version of the LSTM (Cho *et al.*, 2014). Not only GRUs are easier to train than LSTMs, but they also match or outperform LSTMs in terms of accuracy in many tasks (Chung *et al.*, 2014; Tang *et al.*, 2016).

Our model incorporates GRU layers that accept the MHC and the peptide to predict the binding probability. Both the MHC and the peptide are represented as character sequences which let the model predicts even unseen MHCs. The characters were represented as embeddings that encode the information for each amino acid type. Experimental results show that transfer learning from a larger amino acid database can help improve the embeddings, with further improvements possible through the use of additional model fine-tuning. Representing the MHC as sequences instead of different types (one-hot representation) also helps the generalization of the model in most cases. MHCSeqNet compares favorably against NetMHCPan (Jurtz *et al.*, 2017) and MHCflurry (O’Donnell *et al.*, 2018), which also employed deep learning methods. Our final model achieves over 0.99 area under the receiver operating curve (AUC), a significant improvement over other methods, and can provide additional coverage over MHC types not presented in training data.

## METHODS

We trained deep learning models to predict the probability of binding between peptide and MHC allele where a prediction of 0.0 indicates a non-binder and 1.0 indicates a strong binder. Our models accept two inputs: peptide, in the form of amino acid sequence, and MHC allele, in the form of either amino acid sequence or allele name. Figure 1 shows an overview of our deep learning model. The model consists of three main parts, namely the peptide input processing (Figure 1 A and C), the MHC allele input processing (Figure 1 B and D), and the output layer (Figure 1 E). The input processing modules try to learn the best internal representations for the peptide and the MHC. They then pass the processed representations to the output layer which perform the final classification. In the following segments we will go over each part of the model.

**Figure 1.**
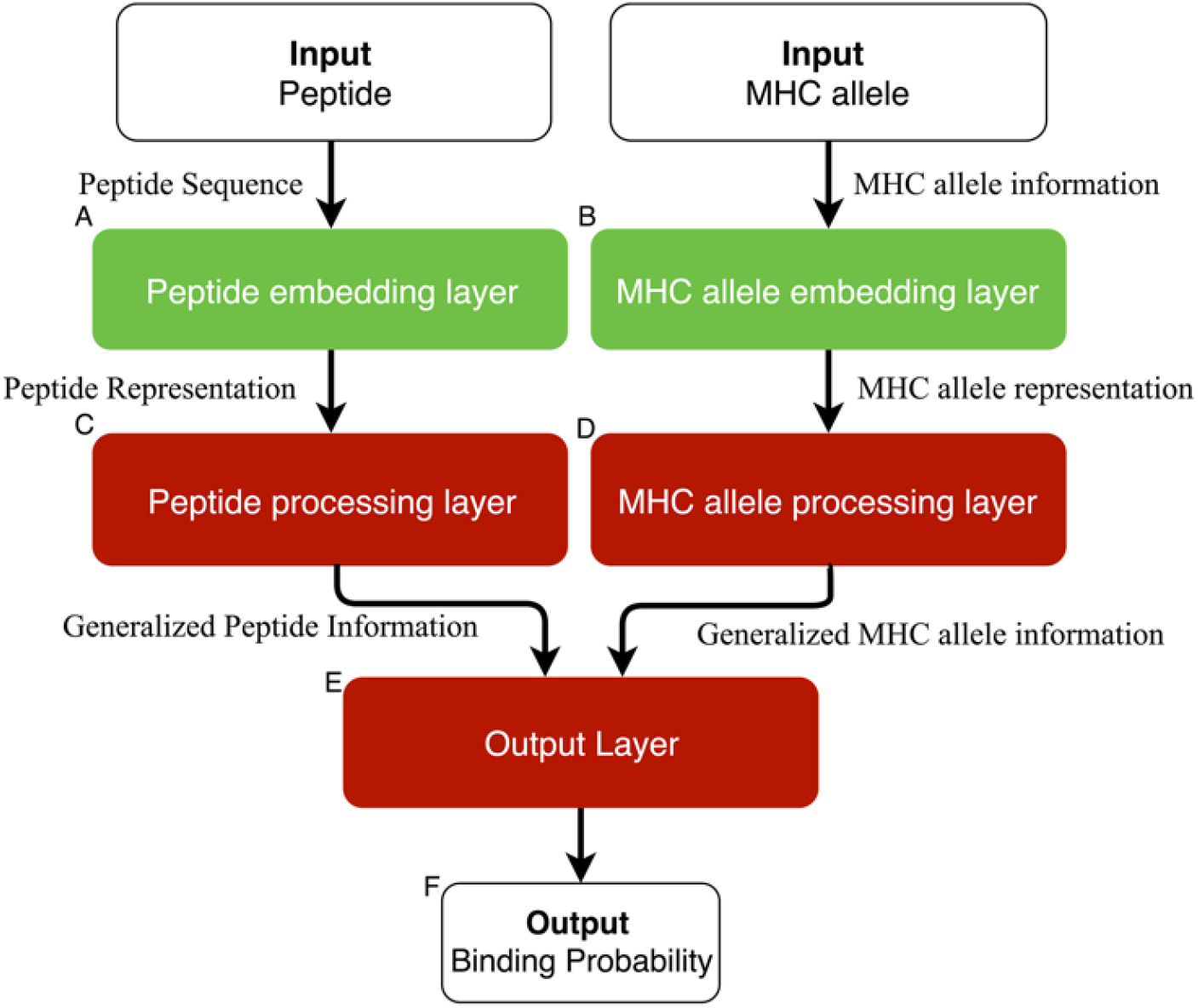
An overview of the MHCSeqNet’s architecture. The model is comprised of three main parts: the peptide sequence processing part (A & C), the MHC processing part (B & D), and the main processing part which accepts the processed information from the previous parts (E). The entire model is a single deep learning model which can be trained altogether.

### Peptide Embedding Layer

We considered two representation models for amino acids. The first is a simple one-hot model where each amino acid is represented by a unit binary vector, e.g. [1,0,0,…] for one amino acid and [0,1,0,…] for another amino acid. The second is a continuous vector representation, called embeddings (Collobert *et al.*, 2011), one of the most successful models in Natural Language Processing (NLP) which can capture the semantic and syntactic relationships between words in a sentence. In our case, a peptide may be considered a sentence and amino acids the individual words. Using embeddings allows us to train the representations on a much larger dataset than the target task (pre-training). The embeddings can then be used or adapted to tasks with smaller datasets.

The Skip-Gram Model (Demeester *et al.*, 2016) was used to train the continuous vectors by treating each set of 1 or 3 consecutive amino acids (1-gram or 3-gram) as a unit. The choice of the 3-gram model, also called ProtVec, was selected according to an earlier study (Asgari and Mofrad, 2015). For each peptide, there are three different 3-gram representations with 0, 1, or 2 amino acids offset from the N-terminus of the peptide sequence. For the tunable parameters, we tested window size of 3, 5, or 7 and embedding dimension of 4, 5, and 6 for the 1-gram model. We found the exact choice of these parameters have little effect on the performance of the model. In the case of the 3-gram model we tested with the same set of window sizes but only at a fixed embedding dimension of 100, which was the reported optimal parameter for this model (Asgari and Mofrad, 2015).

### Peptide Processing Layer

GRU was chosen as the peptide processing layer (see Figure 1C) because it is capable of processing sequences with variable lengths and generalizing relationship between the amino acid representation in peptide sequence. We used one GRU layer. For the 1-Gram amino acid representation, a bi-directional GRU was used. For the ProtVec model (3-gram), three parallel GRU layers (one for each offset) were required. The number of GRU units tested ranged from 32 to 224.

The input to the GRU is the embeddings from the previous section. We conducted an experiment to find the best amino acid representation for the task by using these possible representations to initialize the peptide embedding layer. Then, we used 5-fold cross-validation to train and test the entire model to find the best peptide representation. We also allowed the model to adapt parameters in the peptide embedding layer via back-propagation to improve the entire model. However, the large number of parameters in ProtVec representation caused the model to overfit during the adaptation and ultimately worsened the performance. To overcome this problem, the ProtVec model was trained with the following procedure. First, the model was trained while freezing the embedding vector until the loss steadied. Then, we enabled the adaptation and continued to train the model for 1-3 epochs. Finally, the embedding parameters were frozen again, and the model was trained until it stopped improving. This procedure is similar in spirit to other transfer learning methods used in other applications (Felbo *et al.*, 2017; Grézl *et al.*, 2014).

### MHC Allele Representation

We considered representing an MHC allele with its amino acid sequence in order to allow the model to predict binding affinity values for new MHC alleles that were not part of the training dataset, as long as the new allele’s amino acid sequence is known beforehand. We obtained the amino acid sequence of human MHC class I alleles from the Immuno Polymorphism Database (IPD) (Robinson and Marsh, 2010) in February 2018. Then, we extracted the amino acid sequence portions that correspond to the two alpha helices that participate in the binding with peptide ligand (Jurtz *et al.*, 2017), e.g. the residues 50-84 and 140-179 on the structure of HLA-B*35:01 (PDB ID: 1A1N, Supplementary Figure 1) (Smith *et al.*, 1996). To map these residue locations on each allele, we created an amino acid multiple-sequence alignment of all alleles using MUSCLE v 3.8.31 (Edgar, 2004).

Each MHC allele is inputted into the MHC allele embedding layer as a sequence of amino acids (see Figure 1B). The MHC allele embedding layer was randomly initialized instead of using pre-trained weights because the alignments can contain gaps which were represented with a special character. For the MHC allele processing layer (see Figure 1D), two different layers were tested, namely the fully connected layer and the GRU. We tuned by varying the layer size from 50 to 350 neurons for the fully connected layer, and from 8 to 80 for the GRU. In addition, we conducted another experiment using a one-hot representation of the MHC allele to compare the performance. This last representation ignores the sequence of the MHC allele and therefore captures no relationship between any two MHC alleles. Unlike the other representations, the MHC allele one-hot representation is passed to the output layer directly.

### Output Layer

The outputs from both the peptide processing layer and the MHC allele processing layer are then passed through two fully connected layers with rectified linear unit (ReLU) as the activation function (see Fig. 1E). A final classification layer employs a sigmoid activation function to obtain the final output. We tuned the size of the fully connected layers in the range of 64 to 512. In order to prevent overfitting, dropouts (Srivastava *et al.*, 2014) are also applied at the fully connected and GRU layers (Cho *et al.*, 2014). Dropout layers with probability ranged from 0.0 to 0.5 were tested.

### Neural Network Training

We employed a 5-fold cross-validation scheme where the dataset was divided into five partitions of equal size. For each fold four partitions were used for training and the remaining partition for testing. Furthermore, in each fold, we separated 20% of the training data for hyperparameter tuning and early stopping of the training process. In our model, architecture weights are shared across all MHC alleles, as supposed to building one model per MHC allele (O’Donnell *et al.*, 2018). Adam optimization algorithm was used for the training (Kinga and Adam, 2015).

#### The Final Neural Network Prediction Model

The final model was developed by using the best representation of peptide and MHC alleles as well as the neural network layers to process those inputs. The best representation of peptide is 1-Gram amino acid trained on the combination of Swiss-Prot and Human Protein as described below. For the best representation of the MHC allele, there are two representations which produce similar results. The first representation is the one-hot of sequence of amino acid and the best layer to process this sequence is fully connected layer. The second representation is the one-hot MHC allele. This representation, however, cannot make prediction for new MHC alleles. Thus, we included both models in the software. We then created a simple ensemble model using the model trained on each fold. The final binding prediction is determined by finding the median of the probability output from the 5 models.

### Performance Evaluation

The performance of our models was measured using the area under the receiver operating curve (AUC). The AUC were calculated for both the whole test set and for individual MHC alleles. Performance over all 92 MHC alleles were included in the calculation for the whole test set. For individual alleles we only evaluate on 55 MHC alleles that have at least 30 ligands in total and at least 5 ligands each for positive and negative interactions. Finally, the performance across the five folds from the cross-validation were averaged. Additionally, we also report the F1 score for some experiments. The score is calculated by selecting the threshold that achieves the highest F1 from the receiver operating curve.

We compared the AUCs of our models to those of NetMHCPan 4.0 (Jurtz *et al.*, 2017) and MHCflurry version 1.1.0 (O’Donnell *et al.*, 2018). To ensure a fair comparison, we also retrained MHCflurry using our dataset and measured its performance. The default hyperparamerters for MHCflurry was used. However, as the public version of NetMHCPan could not be re-trained using our dataset, we evaluated its performance as is. In each comparison, only MHC alleles and ligand amino acid sequence lengths that were supported by all software involved were considered. This restricted the evaluation sets to 49 MHC alleles for NetMHCPan and 52 alleles for the original MHCflurry. Furthermore, NetMHCPan supports only ligands between 8 and 11 amino acids long.

Additionally, we tested all models on an external dataset (Bassani-Sternberg *et al.*, 2016) that consists of MHC class I peptidome from four human individuals whose HLA-A, HLA-B, and HLA-C alleles have been determined. Since a detected ligand in this dataset could be bound to any of the MHC alleles present, we took the maximal predicted binding probability over the set of MHC alleles as the predicted binding for each model. To enable the calculation of AUC here, we also include the negative data from the test set described above which were not used during the training of our models in each fold of the cross-validation.

### Prediction for New MHC Alleles

We evaluated the capability of our sequence-based model to predict peptide-MHC binding for new MHC alleles by purposely omitting one MHC allele at a time from the training dataset, retraining the model, and then predicting the binding for peptides contained in the omitted data. This process was repeated for every MHC allele in the dataset and the prediction performance in term of average AUCs over 5 models (derived from 5-fold cross-validation) were recorded. The one-hot model, which conveys no information across MHC alleles, was used as the performance baseline.

## MATERIALS

### Peptide Sequence data

To obtain a large amount of amino acid sequences for pre-training the 1-gram and 3- gram models described above, we downloaded 468,891 verified protein sequences of all species from Swiss-Prot (Bateman *et al.*, 2017) and also constructed a dataset of 16 million simulated proteasome-cleaved human 9-mer peptides using NetChop (Keşmir *et al.*, 2002). Likely 9- mers were defined as those flanked by proteasome cleavage site with predicted cleavage probability ≥ 0.5. The 1-gram and 3-gram models were than trained on the Swiss-Prot proteins alone, the simulated 9-mer peptides alone, or the combination of the two.

### Peptide-MHC binding affinity data

We combined peptide-MHC binding affinity data from IEDB (Robinson and Marsh, 2010) and MHCflurry (O’Donnell *et al.*, 2018) and selected entries corresponding to human MHC class I molecules (HLA-A, HLA-B, and HLA-C) with peptide ligand lengths between 8 to 15 amino acids. Entries with ambiguous ligand amino acids, namely B, X, J, and Z, or nonspecific MHC alleles, such as HLA-A30 or MHC class I, were excluded. Furthermore, as one of our ANN models needs the amino acid sequence of MHC allele as one of the input, only alleles whose sequences are available in the Immuno Polymorphism Database (Robinson and Marsh, 2010) were retained.

We chose to disregard all quantitative binding affinity values and used only qualitative binding classifications, namely, Positive-High, Positive, Positive-Intermediate, Positive-Low, and Negative, because the binding affinity values were acquired through diverse experimental techniques performed in multiple laboratories. Furthermore, we found that removing low-confidence entries (Positive-Intermediate and Positive-Low) slightly improved the prediction performances. There were also a number of conflicting entries which contained the same MHC allele and ligand but opposite binding classification. As the majority of conflicts came from a few sources, we decided to exclude all entries from them. For each of the remaining conflicts, we reassigned the binding classification based on a majority vote. If there is no majority, all of the associated conflicting entries were excluded. Lastly, duplicated entries were discarded.

In total, the final dataset contains 228,348 peptide-MHC entries consisting of 31 HLA-A, 49 HLA-B, and 12 HLA-C alleles. The number of ligands per allele ranges from 41 to 21,480 (Supplementary Data).

## RESULTS

### MHC Allele Representation

We trained our models using two different MHC allele representations: a one-hot system which conveys no relationship between MHC alleles, and an amino acid sequence-based representation that permits ligand inference across alleles. We also tested two entry layer architectures for reading the MHC allele sequences, a fully connected layer and a GRU layer (see Methods). Evaluations based on AUC showed that using a fully connected layer as the entry layer gives better performance than using a GRU layer overall (Table. 1) and on 76% of the alleles tested (44 out of 58). Although the one-hot representation yielded the highest overall performance, it outperformed the sequence-based representation with fully connected layer on only 45% of the alleles tested (26 out of 58). A closer inspection revealed that the performance of both models on alleles with a large amount of both positive and negative data were similar. However, on alleles that had less than 25 data points for either the positive or negative class, one-hot based model tended to perform better. This is due to the fact that sequence-based model needs to process the MHC allele sequence, so it requires more parameters and more data to train than one-hot based model.

**Table 1.** Evaluation of MHC allele representation and entry layer models

| Allele Representation | Sequence-Based |  | One-Hot |
| --- | --- | --- | --- |
| Entry Layer Model | Fully Connected | GRU | None |
| Overall AUC<sup>a</sup> | 0.9910 | 0.9898 | 0.9917 |

<sup>a</sup>Values shown here were obtained from five-fold cross-validation

### Peptide Embedding Layer

The one-hot, 1-gram, and 3-gram peptide representations were pre-trained on three different datasets: Swiss-Prot proteins (“Swiss-Prot”), simulated human proteasome-cleaved 9-mers (“Human”), and the combination of the two (“Combined”). We also studied the case where no pre-training is done, and the embedding layer was initialized randomly (“None”).

Overall, the 1-gram model yielded the best performance, with an average AUC of 0.991715, followed by the one-hot model, with an average AUC of 0.991680 (Table 2). For the 1-gram model, combining the two amino acid sequence datasets improved the performance over using individual dataset and using no dataset. Rather unexpectedly, regardless of the adaptation method (see Methods) or the dataset tested, the 3-gram model, which has previously been used to analyzed protein structural family (Asgari and Mofrad, 2015), was consistently outperformed by the one-hot and 1-gram models. We suspected that the large number of parameters in the 3- gram model caused the model to be overfitted even when a special adaptation method (“Partial” in Table 2) is used. Nonetheless, significant improvement in AUC was achieved with our improved adaptation method.

**Table 2.** Comparison of peptide representation models and adaptation methods

| Representation Model | Sequence Dataset | Adaptation | Overall AUC |
| --- | --- | --- | --- |
| **None** | None | No | **0.991680** |
| **1-Gram** | Swiss-Prot | Full | 0.991401 |
|  | Human | Full | 0.991136 |
|  | Combined | Full | **0.991715<sup>a</sup>** |
|  | None | Full | 0.990859 |
| **3-Gram** | Swiss-Prot | No | 0.956362 |
|  | Human | No | 0.96069 |
|  | Combined | No | 0.959503 |
|  | None | No | 0.957562 |
|  | Swiss-Prot | Full | 0.967181 |
|  | Human | Full | 0.96637 |
|  | Combined | Full | 0.966889 |
|  | None | Full | 0.966244 |
|  | Swiss-Prot | Partial | **0.982490** |
|  | Human | Partial | 0.98119 |
|  | Combined | Partial | 0.981133 |
|  | None | Partial | 0.981052 |

<sup>a</sup>The 1-gram model trained on combined dataset was selected for the final model

### Evaluation Using MHC Class I Binding Dataset

From above results, we determined the best peptide representation to be the 1-gram model trained on the combination of Swiss-Prot proteins and simulated human proteasome-cleaved 9-mers. For the MHC representation, because the one-hot model performed better given sufficient amount of data while the sequence-based model performed better when the amount of data was limited, we evaluated both models here. Our models were then compared to NetMHCPan version 4.0 (Jurtz *et al.*, 2017), MHCflurry version 1.1.0 (O’Donnell *et al.*, 2018) and the retrained MHCflurry, using a five-fold cross-validation scheme (see Methods). This revealed that all of our models significantly outperformed both NetMHCPan and MHCflurry overall (Figure 2) and on 77%-84% of the individual MHC alleles tested in terms of AUC. Compared to NetMHCPan, our one-hot and sequence-based MHC representation models achieved better AUC on 43 and 40 out of 49 alleles supported by both software. Compared to the already trained MHCflurry, both of our models performed better on 40 out of 52 alleles. Finally, although the overall AUC for MHCflurry improved considerably after re-training with our dataset, both of our models still achieved better AUC in 49 out of 58 alleles. Additionally, we calculated F1 score on the whole test set at the threshold which yielded the highest F1 score for each model. The results show that all of our models are still the best in this regard.

**Figure 2.**
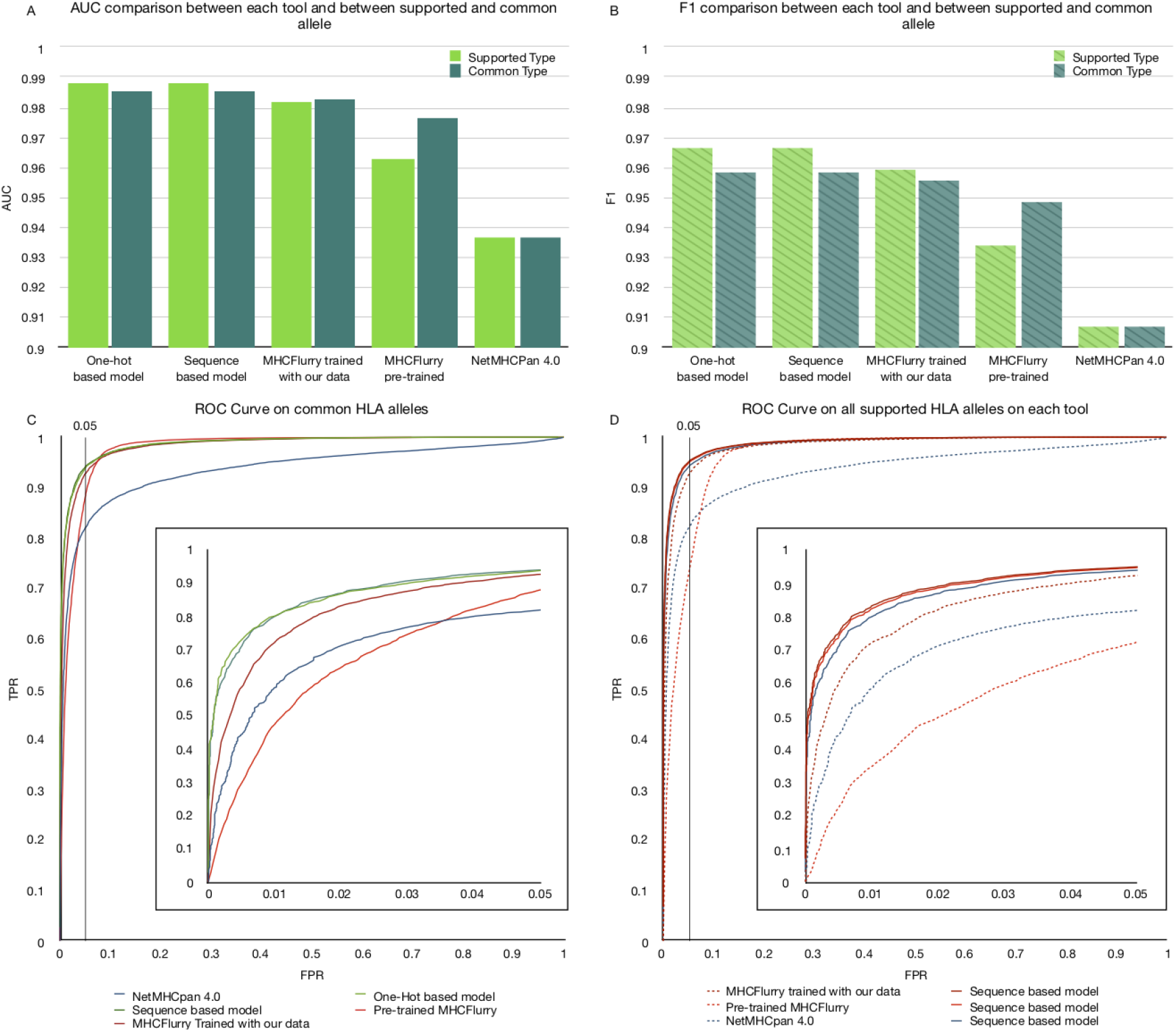
MHCSeqNet achieves the best AUC and F1 values on MHC class I binding dataset. (A) Bar plots showing the AUC values of each tool when evaluated on the set of all MHC alleles it supported (Supported Type) and on the set of MHC alleles supported by all tools (Common Type). (B) Similar bar plots for F1 values. (C) The ROC plot for all tools when evaluated on the set of MHC alleles supported by all tools. Vertical black line indicates the 5% FDR. Inset shows the zoomed in ROC plot for the region with ≤5% FDR. (D) Similar ROC plot for the evaluation on MHC alleles supported by individual tools.

### Evaluation Using MHC Class I Peptidomes

We further evaluated our models, NetMHCPan and MHCflurry on a MHC class I peptidome dataset (Bassani-Sternberg *et al.*, 2016) which were derived from mass spectrometry analyses of four human individuals whose HLA-A, HLA-B, and HLA-C alleles have been typed (Table 3). Because it is unclear which MHC allele was bound to each detected ligand, the maximal predicted binding probability over the set of MHC alleles in each sample was designated as the final predicted binding affinity for each model. It should be noted that some HLA-C alleles were not supported by NetMHCPan, MHCflurry, or our one-hot model (Table 3), and that the sequence-based model alone could make predictions for these alleles. Both of our models achieved the best overall AUC and F1 score in all cases (Figure 3). As observed earlier, sometimes our model that uses the one-hot MHC allele representations yielded better performance, and sometimes our sequence-based model did, depending on the number of training data points for the MHC alleles involved (Supplementary Table 1).

**Table 3.** Typed MHC alleles of four individuals from the HLA class I peptidome dataset

| Sample ID | Mel 12 | Mel 15 | Mel 16 | Mel 8 |
| --- | --- | --- | --- | --- |
| **HLA-A** | A*01:01 | A*03:01 | A*01:01 | A*01:01 |
|  | - | A*68:01 | A*24:02 | A*03:01 |
| **HLA-B** | B*08:01 | B*27:05 | B*07:02 | B*07:02 |
|  | - | B*35:03 | B*08:01 | B*08:01 |
| **HLA-C** | C*07:01<sup>a</sup> | C*02:02<sup>b</sup> | C*07:01<sup>a</sup> | C*07:01<sup>a</sup> |
|  | - | C*04:01 | C*07:02 | C*07:01<sup>a</sup> |

<sup>a</sup>Alleles not supported by the original MHCflurry
<sup>b</sup>Allele not supported by MHCflurry, NetMHCpan, and our one-hot model

**Figure 3.**
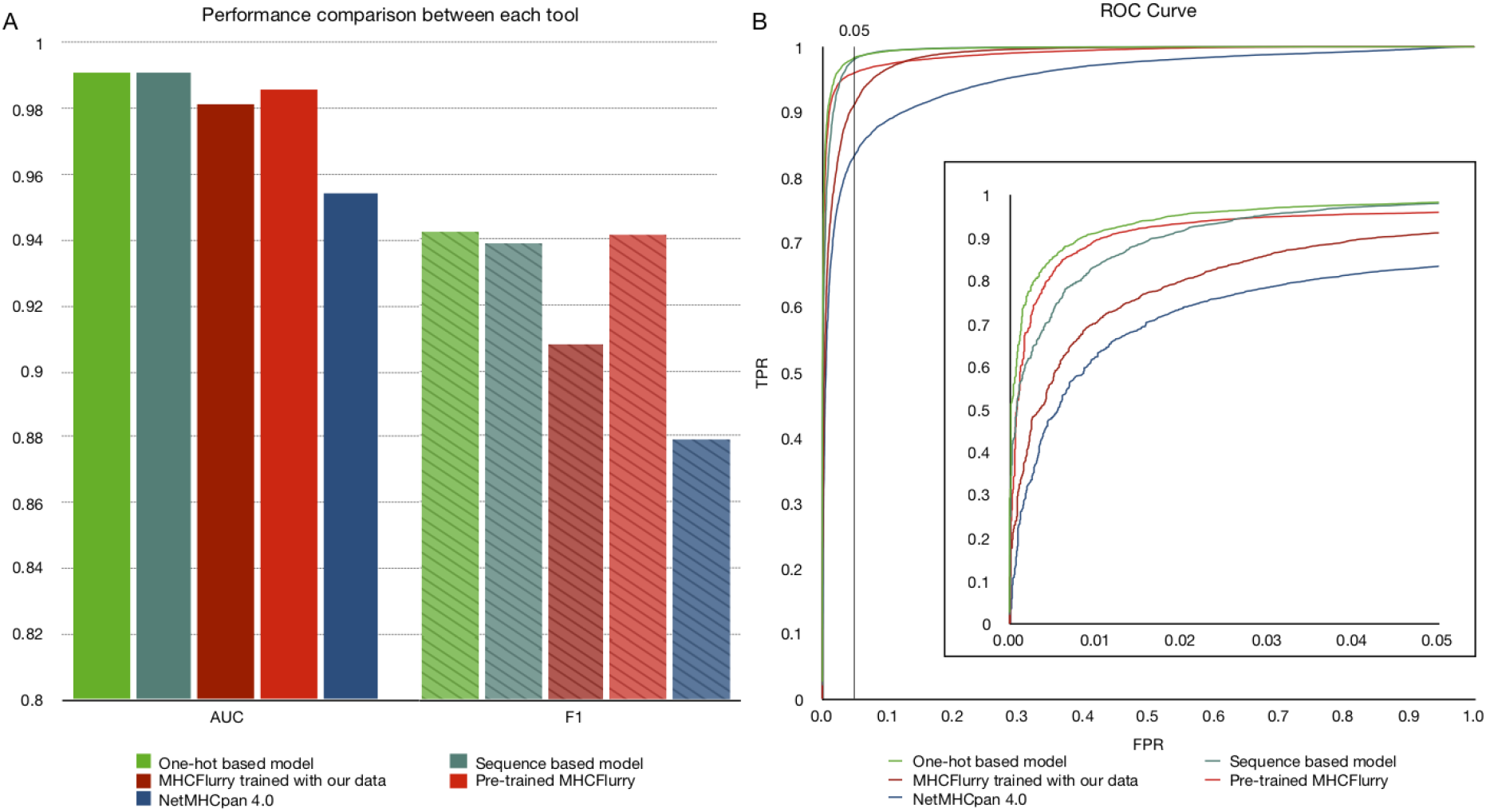
MHCSeqNet achieves the best AUC and F1 values on MHC class I peptidome dataset. (A) Bar plots showing the AUC and F1 values of each tool. (B) The ROC plot for all tools. Vertical black line indicates the 5% FDR. Inset show the zoomed in ROC plot for the region with ≤5% FDR.

### Prediction for New MHC Alleles

To evaluate our model’s ability to predict ligand binding probability for new MHC alleles when MHC alleles are represented by their amino acid sequences, we trained the model using data from all-but-one alleles and used the omitted allele for the testing. To ensure the correctness of the results, we performed the experiment 5 times with different random initializations and calculate the average AUC and the Standard Deviation (SD) for each held-out allele. Compared to the baseline one-hot MHC allele representation model, the AUC improved for 59 out of 65 MHC alleles tested, with median increasing from 66.34% to 89.12% (Supplementary Figure 2). Additionally, the SD values averaged across held-out alleles for the one-hot and sequence-based models are 5.30% and 1.63%, respectively. Thus, the sequence-based model is more accurate and more stable than one-hot-based model overall.

## DISCUSSION

### Impact of Data Cleaning

In addition to removing duplicated entries and entries with ambiguous peptide sequence or MHC allele name as is regularly performed in other studies (Jurtz *et al.*, 2017; O’Donnell *et al.*, 2018), we also examined the impact of conflicting entries (entries with the same peptide sequence and same MHC allele but opposite binding affinity classification) and low-confidence entries (Positive-Intermediate and Positive Low) on the prediction performance. Although conflicting entries constitute less than 5% of the raw dataset, the vast majority of them (15,305 out of 17,914 conflicts) involve the same source, which reported HLA-B*27 ligands identified in transgenic mice (Barnea *et al.*, 2017), and should be entirely excluded or at least carefully scrutinized. The remaining conflicts could be resolved by majority voting which slightly improves the prediction performance. We also found that the exclusion of low-confidence entries slightly improved the prediction performances when tested on all entries or only on high-confidence entries.

### Performance on MHC class II

Our model has the potential to predict ligand for MHC class II alleles as it can accept peptides of any length. We tested the one-hot model on 22 MHC class II alleles with at least 150 entries in the binding affinity dataset (at least 5 must be positive and at least 5 must be negative), and compared its performance to MHCflurry, which support peptides of up to 15 amino acids. Our model performed slightly better for peptides of up to 15 amino acids (average AUC of 0.83792 compared to 0.82147, 13 out of 22 alleles, Supplementary Table 2) and achieved moderate accuracy for longer peptides (average AUC of 0.63878). It should be noted that the performance evaluation on longer peptides here likely suffered from the lack of data, especially the lack of the negative entries, because it was based on only seven alleles, five of which contain less than 100 entries and less than 30 negative entries. When the predictions for all these seven alleles were pooled together, the AUC improved significantly to 0.9823.

### Capability to make prediction for new MHC alleles

The capability to predict binding affinity of a candidate neoepitope against all MHC alleles presented in a patient is highly desirable because neoepitopes that can bind to multiple MHC alleles are likely to be immunogenic. The sequence-based version of MHCSeqNet not only better predicted ligands for new MHC alleles than its one-hot counterpart (Supplementary Figure 2) but also improved over existing tools (Figure 2 and 3). Curiously, the sequence-based model did not outperform the one-hot model when tested on a peptidome dataset which contains a few HLA-C alleles not present in the training dataset (Table 3, Supplementary Table 1). We suspected that this is due to the overall lack of HLA-C data (HLA-C entries constitute only 3% of the final training dataset) for the sequence-based model to learn from. More data on HLA-C epitopes from future experiments should further improve the performance of the sequence-based version of MHCSeqNet. In light of its flexibility of usage and competitive performance, MHCSeqNet should contribute strongly to the screening of effective neoepitopes for cancer vaccine development.

## ACKNOWLEDGEMENTS

This work was partly supported by the Ratchadapisek Sompoch Endowment Fund at Chulalongkorn University [Grant for the Establishment of Research Group to S.S. and E.C.; Grant for the Development of New Faculty Staff to N.P.] and the Thailand Research Fund [MRG6080087 to N.P.].

## CONTRIBUTIONS

P.P., N.P., E.C., and S.S. analyzed data and wrote the manuscript. P.P. developed the MHCSeqNet software tool. N.P., E.C., and S.S. directed and supervised the project.

## Conflict of interest statement

None declared.

